# Processing the diffusion-weighted magnetic resonance imaging of the PING dataset

**DOI:** 10.1101/2020.11.24.396549

**Authors:** Noor B Al-Sharif, Etienne St-Onge, Guillaume Theaud, Alan C Evans, Maxime Descoteaux

## Abstract

Diffusion-weighted magnetic resonance imaging (dMRI) allows for the in-vivo assessment of anatomical white matter in the brain, thus allowing the depiction of structural connectivity. Using structural processing techniques and related methods, a growing body of literature has illustrated that connectomics is a crucial aspect to assessing the brain in health and disease. The Pediatric Imaging Neurocognition and Genetics (PING) dataset was collected and released openly to contribute to the assessment of typical brain development in a pediatric sample. This current work details the processing of diffusion-weighted images from the PING dataset, including rigorous quality assessment and fine-tuning of parameters at every step, to increase the accessibility of these data for connectomic analysis. This processing provides state-of-the-art diffusion measures, both classical diffusion tensor imaging (DTI) and more advanced HARDI-based metrics, enabling the evaluation not only of structural white matter but also of integrated multimodal analyses, i.e. combining structural information from dMRI with functional or gray matter analyses.

## 1. Introduction

### Brain structure & development

Post-natal neural development is characterized by widespread structural changes of the brain, in both gray and white matter regions [Sowell et al., 2004; Lenroot and Giedd, 2006; Giedd and Rapoport, 2010]. Microstructural changes in brain architecture, such as cortical thickness or myelination, lead to global morphometric changes detectable by magnetic resonance imaging (MRI) techniques, including anatomical images (such as T1 or T2^*\**^ weighted) and diffusion-weighted imaging (dMRI). Just as these techniques can be used to assess differences between health and disease states, they also provide insight into underlying microstructural changes in the brain that characterize typical development *in vivo* [Lim and Helpern, 2002; Prayer and Prayer, 2003; Qiu et al., 2015]. Implementing multimodal imaging methods in the assessment of structural development throughout the lifespan enables a comprehensive perspective of anatomical growth and variance across the whole brain. Research has shown that a major component of post-natal brain development occurs in the strengthening or refining of local and whole-brain white matter connections [Lebel et al., 2008; Jernigan et al., 2011]. Many of these changes are experience-dependent and causally linked with myelination, synaptogenesis and synaptic pruning in different areas of the brain at different developmental periods [Shatz, 1990; Benes et al., 1994; Paus et al., 1999; Stiles and Jernigan, 2010; Deoni et al., 2011]. Such changes in structural white matter provide a compelling case for the use of dMRI in assessing brain development.

### Diffusion MRI & white matter processing

DMRI is a particular MRI sequence that quantifies water displacement in multiple directions. From diffusion-weighted images, local diffusion models are reconstructed to estimate the underlying characteristics of white matter anatomy [Jones, 2008; Descoteaux, 2015]. The dynamic changes of this microstructure in development can be characterized both in structural and functional analyses [Klingberg et al., 1999; Mukherjee et al., 2002; Olesen et al., 2003; Sotiropoulos and Zalesky, 2019]. These changes, whether localized or not, underlie global changes in whole-brain networks, which form the basis of brain function and behaviour [Nagy et al., 2004]. As such, it is important to evaluate these changes in the context of brain connectivity, using techniques such as dMRI to depict structural white matter changes [Jeurissen et al., 2019]. As of yet, we have a limited understanding of how white matter develops during childhood and adolescence and therefore require more insight into how these structures change over time.

Though the implementation of dMRI techniques has improved our understanding of anatomical white matter structure, processing of such data is known to be rife with complex challenges [Jones and Cercignani, 2010]. These challenges consist of low spatial resolution, noisy acquisition, MRI gradient-related artifacts and fiber organisation ambiguity (crossing, kissing, fanning) [Jbabdi et al., 2015; Dell’Acqua and Tournier, 2019; Jeurissen et al., 2019]. As such, processing dMRI data needs to be reliable and reproducible to improve dMRI quality and better depict local and global structure of the human brain [Theaud et al., 2020].

### PING

In order to depict stereotypical changes or patterns in structural development, the use of large-scale population-representative datasets is essential. One such dataset is the Pediatric Imaging Neurocognition and Genetics (PING) study: a large-scale developmental cohort of subjects aged between 3 and 22 years, including imaging, cognitive and genetic data. This comprehensive study provides a rich multimodal imaging perspective of development, from childhood to early adulthood, and enables the integration of behavioural and genetic information. PING publicly provides both the raw imaging and demographic data, as well as some processed derivatives of these data including cortical thickness, surface area, volumes and some diffusion-tensor measures. However, despite this notable amount of data made available by PING, structural white matter connectivity measures were not included, to our knowledge.

To best take advantage of these large-scale datasets, such data must be made accessible for use beyond their originating research groups. These types of open science practices promote scientific collaboration and discovery, such as the Human Connectome Project (HCP) [Glasser et al., 2013]. The creation and analysis of large-scale population datasets are arduous tasks best accomplished as collaborative efforts. Sharing data encourages better, more transparent research practices and also minimizes the necessity for repetition in laborious data collection and processing. Therefore, the derivatives of this current processing will be made publicly available and accessible through the National Database for Autism Research (NDAR) repository, a database of the National Institute of Mental Health Data Archive (NDA), which also hosts the original PING dataset.

## 2. Methods

In this section, we specify how the initial PING release and dMRI acquisition were harmonized and formatted [Jernigan et al., 2016]. The pre-processing and processing applied to dMRIs is detailed, along with any processing applied in parallel to the T1-weighted anatomical images. Finally, tractography and connectivity matrix generation are described. As detailed in Jernigan et al. [2016], written consent was obtained from parental guardians for participants below 18 years of age and from the participants themselves otherwise, in accordance with IRB protocols. For reproduction purposes, details are listed in the Usage Notes and Code availability sections. Figure 1 illustrates a comprehensive overview of the processing pipeline.

**Figure 1:**
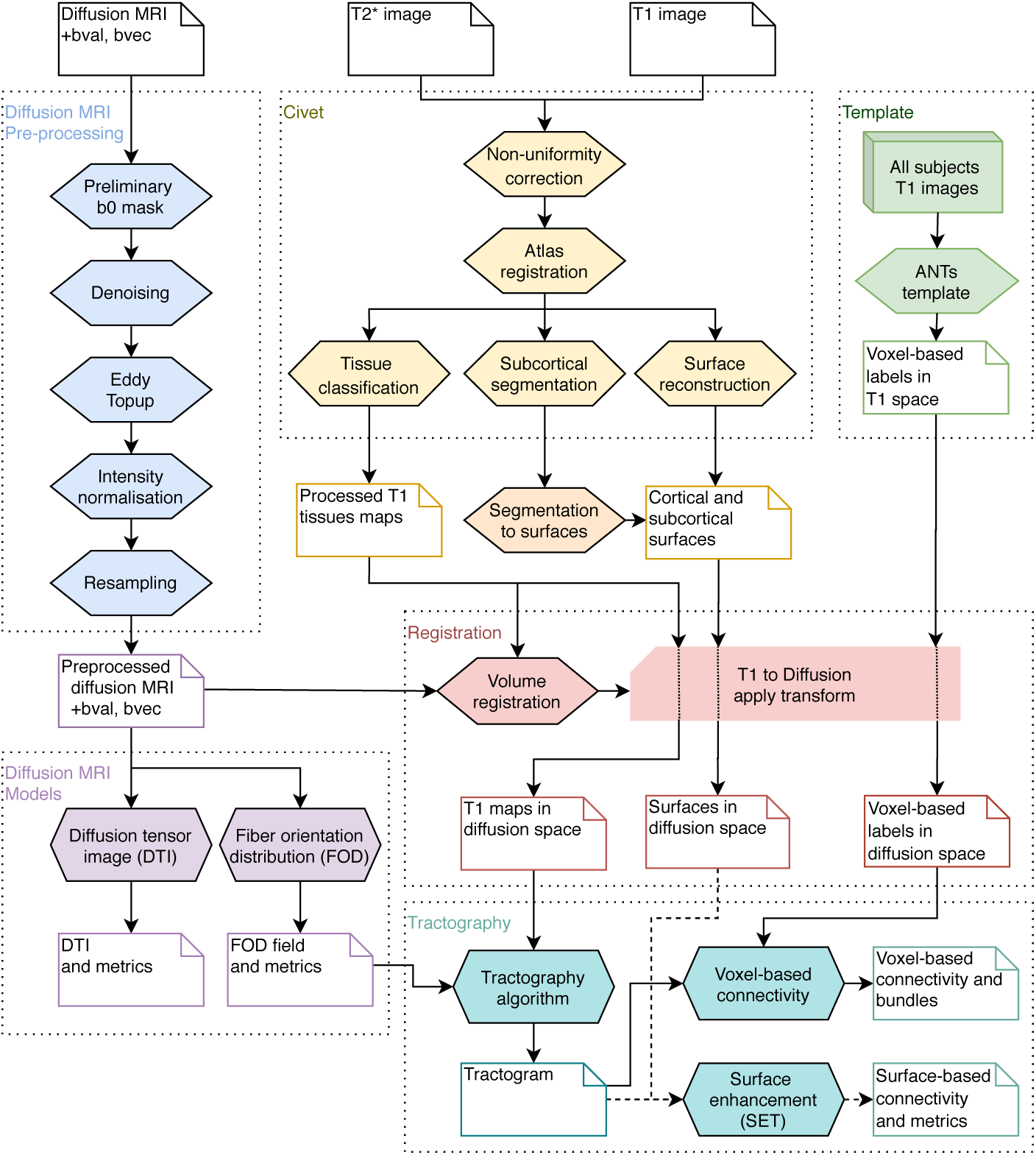
Processing pipeline architecture, for diffusion, T1 and T2^*\**^ weighted images: dMRI pre-processing (blue), CIVET for WM-GM-CSF tissue maps with surfaces (yellow), T1 template reconstruction (green), Registration to align T1 maps from anatomical to diffusion space (red), diffusion local models reconstruction (purple), and tractography with structural connectivity (cyan).

### 2.1. Input data collection and formatting

#### Imaging data collection

Original PING data were collected in Digital Imaging and Communications in Medicine (DICOM) format across nine sites on four 3T MR machine types: Siemens TrioTim, GE Discovery, GE Signa, and Philips Achieva. Scanning protocols were harmonized across all scanners and sites to minimize variance across scans. From PING, we gained access to a total of 954 subjects with dMRI scans, 634 of which also have at least 2 or more repeated scans within the same session. In total, 1669 unique acquisitions were used as input for the subsequent processing.

#### Anatomical images

Though the PING dataset contains both T1w and T2^*\**^w images, only T1w images are required for the processing described here. T1w images were collected using a real-time prospective motion correction (3D PROMO) technique to minimize the effects of motion, which is typically widespread in pediatric samples. Subjects were processed only if they had both dMRI and T1w images which passed quality assurance, i.e. minimal motion, no major artifacts, etc.

#### Data conversion: DICOM to NIfTI

Since most dMRI processing tools cannot directly input DICOM files, a conversion is necessary. First, DICOM (`‘.dcm’`) files were converted to Neuroimaging Informatics Technology Initiative (NIfTI) file format (`‘.nii’`) using *dcm2niix* [Li et al., 2016]. For dMRI volumes, this conversion also generates two accompanying `‘bval’` and `‘bvec’` files which contain acquisition details of the b-values and b-vectors for each volume.

#### Transformation (space) convention

To facilitate the use of common dMRI software, all NIfTI files were modified to the RAS (left-to-right, posterior-to-anterior, inferior-to-superior) convention.

### 2.2. Diffusion-weighted images (dMRI)

Each dMRI acquisition is composed of a total of 32 diffusion-weighted three-dimensional volumes: 30 with diffusion directions and 2 baseline (b0 images). Of the two b0 images, one is acquired in the anterior-to-posterior (A-P) direction, which is in alignment with all volumes of the main dMRI (named `‘b0.nii’`). The second b0 is acquired in the reverse direction, posterior-to-anterior (P-A), (named `‘b0_rev.nii’`). This reverse b0 is used to correct for susceptibility-induced distortion in dMRI. Each direction measures the signal loss along the chosen orientation (b-vector) with a single b-value of 1000.

### 2.3. Diffusion MRI Pre-processing

A preliminary b0 mask (`‘bet_prelim_b0.nii’`) is estimated with *FSL-bet* from the `‘b0.nii’` dilated with *MRtrix-mask filter*. This is done to mask voxels determined to be outside of the brain tissue and to remove unnecessary computation of invalid voxels. Denoising was performed for all dMRI directions (`‘dwi.nii’`) inside the preliminary mask (`‘bet_prelim_b0.nii’`) with *MRtrix-dMRI denoise*[Tournier et al., 2019].Topup correction map was computed with *FSL-topup*, to correct for common dMRI acquisition deformation, using both oppositely-acquired b0 images (`‘b0.nii’`, `‘b0_rev.nii’`) and applied to all dMRI directions (`‘dwi.nii’`). Eddy current correction was also applied using *FSL-eddy. FSL-eddy* applies both eddy current and distortion correction (generated from *FSL-topup*) to all diffusion volumes; it also returns distortion-corrected `‘bval’` and `‘bvec’` files. A new brain mask (`‘bet_b0.nii’`) is computed after the deformation correction, minimally dilated with *MRtrix-mask filter*. Intensity normalization was employed to establish spatial uniformity of the signal intensity across the dMRI volume using *ANTs-N4BiasFieldCorrection*. Additionally, *MRtrix-dMRI normalise* was used to harmonize the diffusion signal across subjects [Tournier et al., 2019]. Each diffusion volume was resampled with linear interpolation to an isotropic resolution of 1*mm* [Lehmann et al., 1999; Dyrby et al., 2014].

### 2.3. Diffusion MRI models

#### DTI. Dipy-TensorModel

was used to estimate the Diffusion Tensor Image (DTI) [Basser et al., 1994] at every spatial position (voxel), with a weighted least squares method. Subsequently, multiple DTI measures were computed from reconstructed tensors (Figure 2), including the Fractional Anisotropy (FA), Mean Diffusivity (MD), Apparent Diffusion Coefficient (ADC), Radial Diffusivity (RD), and red-green-blue (RGB) map colored in the x-y-z orientation (Figure 3).

**Figure 2:**
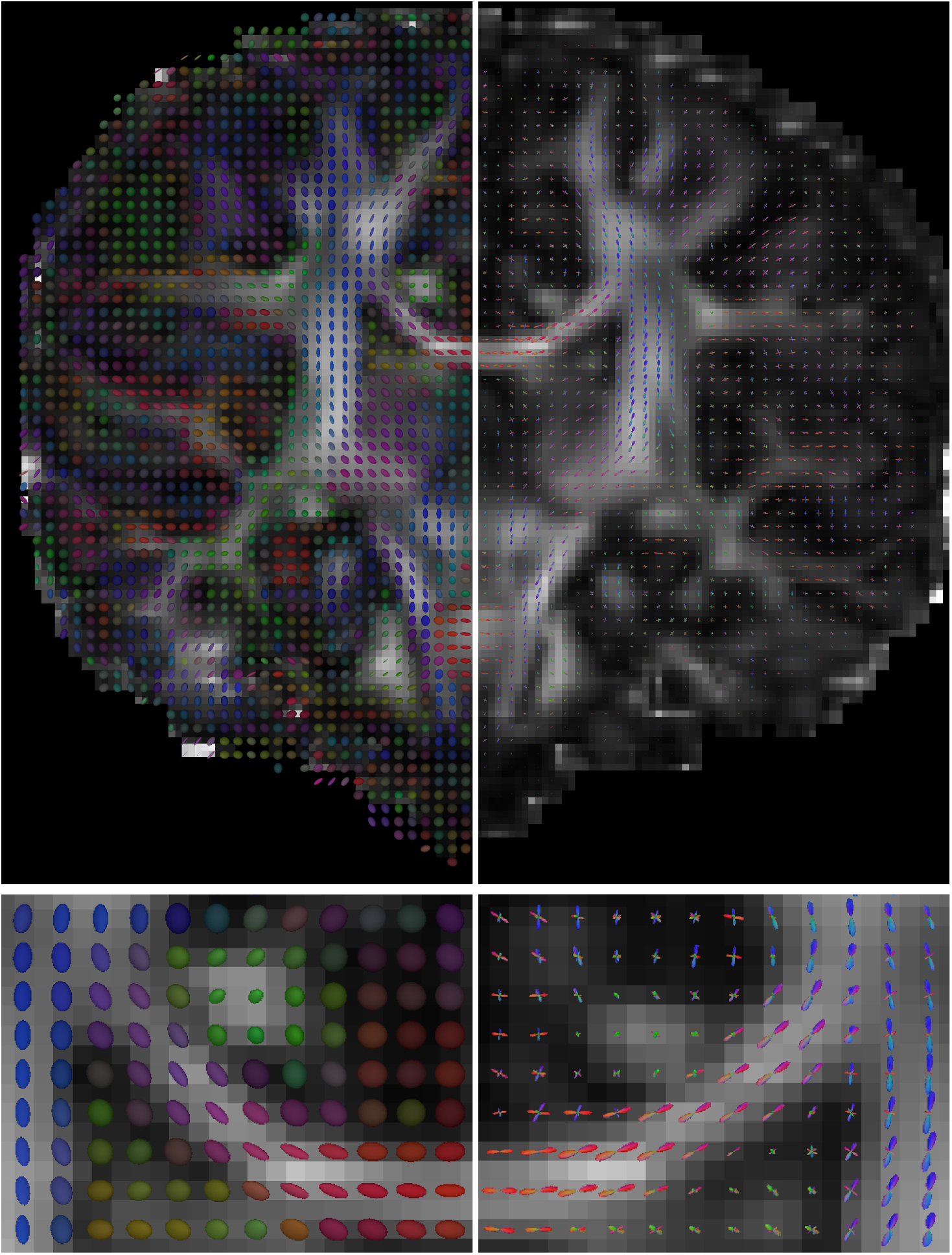
Coronal view of the reconstructed diffusion tensor image (DTI) on the left, and fiber orientation distribution (FOD) on the right. On the bottom row, a zoomed view of the *Corpus Callosum* and *Cingulum* regions.

**Figure 3:**
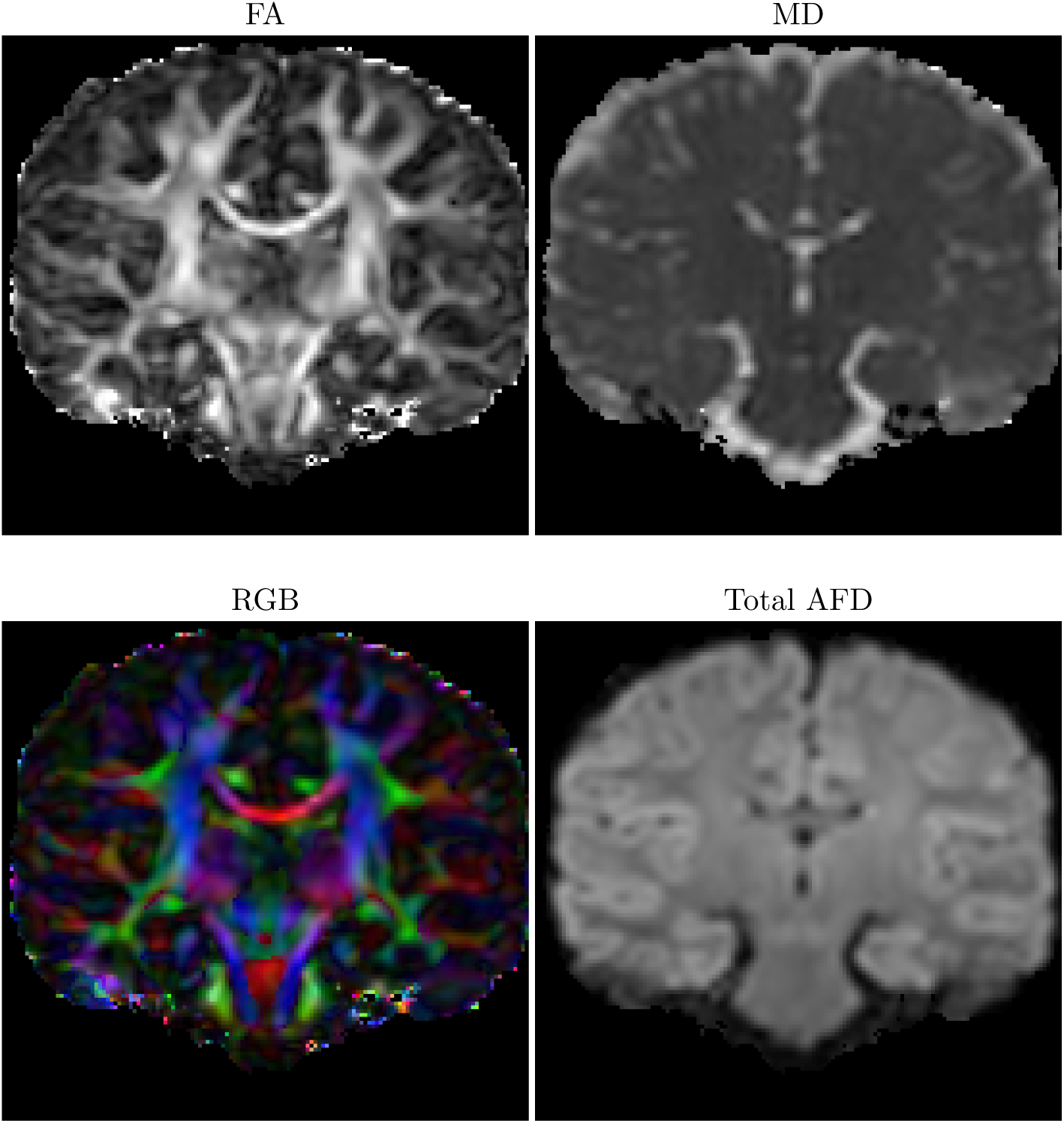
Diffusion tensor image (DTI) field measures: fractional anisotropy (FA), mean diffusivity (MD), and red-green-blue (RGB) map colored from the x-y-z orientation. Total apparent fiber density (AFD) estimated from the Fiber orientation distribution (FOD).

#### FOD. Dipy-csdeconv

package was used to compute fiber orientation distribution (FOD). Initially the “single fiber distribution” is estimated with the `auto response` function and afterward the full distribution is computed at each position using `ConstrainedSphericalDeconvModel` with the `symmetric724` discretized sphere. The FOD represents the estimated orientation distribution of fibrous structure at each voxel [Tournier et al., 2007; Descoteaux et al., 2009]. Peaks, representing main diffusion directions, are extracted from local maxima of each FOD angular distribution, similar to a vector field (Figure 2). This FOD field is later used to compute the tractography and estimate the structural connectivity. In addition, some feature maps are estimated from the FOD reconstruction, illustrated in Figure 3, such as total apparent fiber density (AFD).

### 2.4. T1w image processing

Segmentation maps for white matter, gray matter, cerebrospinal fluid (CSF) and subcortical structures (SC) were generated by *CIVET 2*.*1*, from the native T1 anatomical image. A brain mask, also estimated from this segmentation, was applied to all T1w images before registration [Zijdenbos et al., 1998; Tohka et al., 2004]. Cortical surfaces were reconstructed with *CIVET 2*.*1* [Kim et al., 2005; Lyttelton et al., 2007], similarly to *FreeSurfer*. Subcortical regions were estimated with *ANIMAL* [Collins et al., 1999] and converted into meshes using a marching cube algorithm, and then combined with cortical surfaces in a single file. The anatomical T1 and diffusion volume were co-registered, transforming the T1 volume into native diffusion space. The resulting transformations (affine matrix and warp) from *antsRegistration*, are also used to align subsequent anatomical maps and cortical surfaces to diffusion space [Avants et al., 2011; Theaud et al., 2020].

#### Template construction

In parallel to the included T1w processing within *Tractoflow*, a PING specific T1w template was generated using *antsMultivariateTemplateConstruction2*.*sh* [Avants et al., 2011], which automates an iterative registration process to move any number of T1w volumes into a common space. Once the template has been generated, the resulting transformation files are used to register other template-aligned volumes into individual subject space. This is applied in subsequent processes which require aligning atlas segmentation labels into native T1w and diffusion volume space, as described below in creating voxel-based connectivity matrices. Averaged tissue maps in the template space can be oberved in Figure 4.

**Figure 4:**
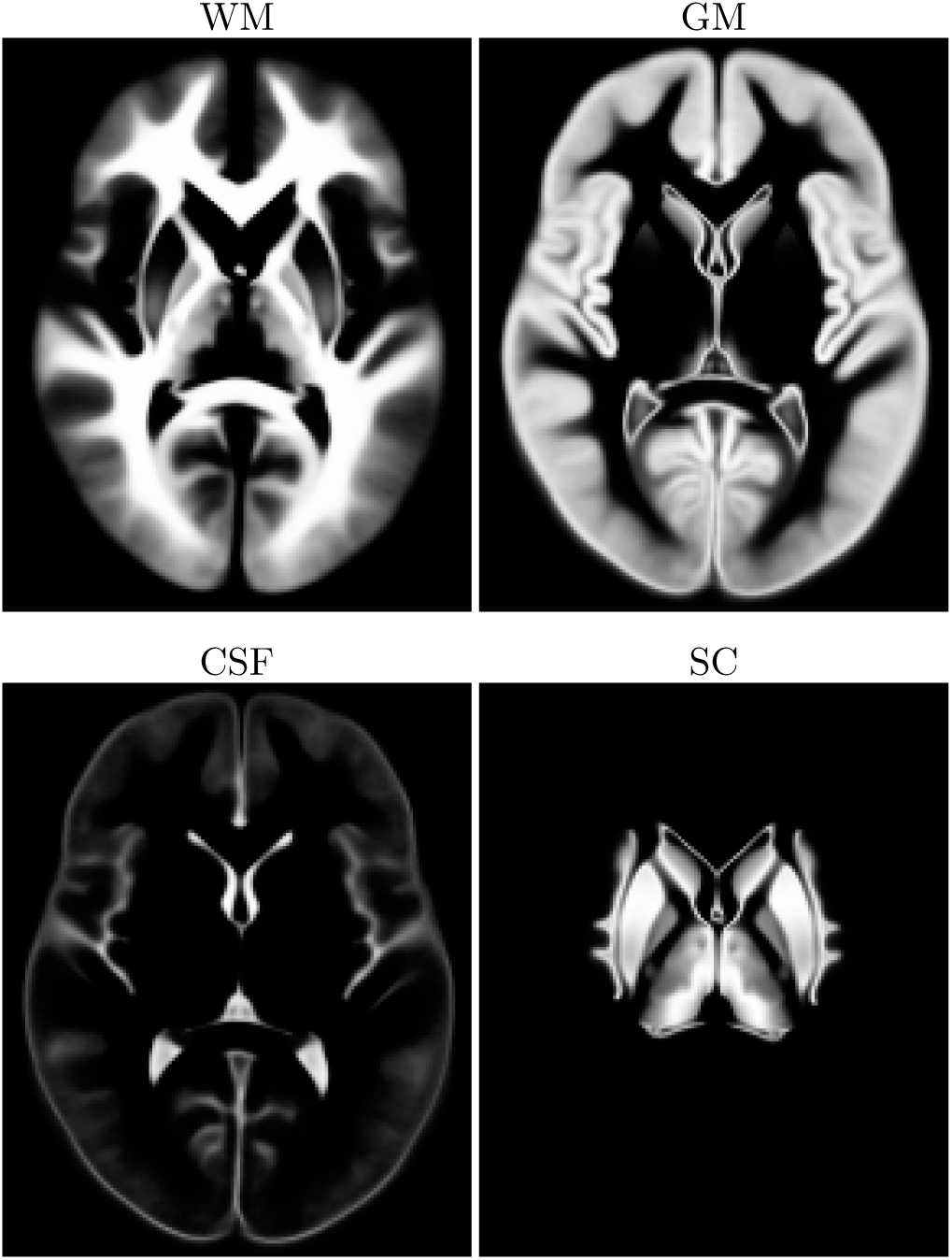
Averaged tissue maps in the template space, axial view.

### 2.5. Structural connectivity analysis

#### Tractography

Structural pathways can be reconstructed from dMRI by following the local orientation of estimated local models with a process called tractography. In this work, we used the “particle filtering tractography” (PFT) [Girard et al., 2014], implemented in *Dipy*, taking advantage of the sharp orientation from the FOD field. Resulting streamlines represent estimated pathways of the brain macrostructure and connectivity. Similarly to anatomically-constrained tractography (ACT) [Smith et al., 2012], PFT takes advantage of previously computed WM-GM-CSF tissue maps, to define legal and illegal regions.

The “surface-enhanced tractography” (SET) [St-Onge et al., 2018], an adaptation of the PFT algorithm, was also used to initialize and terminate streamlines from cortical and subcortical surfaces at the WM-GM boundary. Since CIVET 2.1 produces a standardized mesh, with a fixed number of vertices and triangles for every subject, SET allows for better comparison and individually variable correlation of connectivity features.

#### Connectivity matrices

Connectivity matrices were generated using two separate methods: voxel- and surface-based. Voxel-based matrices were created using a 3D anatomical Desikan-Killiany-Tourville (DKT) [Desikan et al., 2006; Klein and Tourville, 2012] segmentation label in each acquisition’s native diffusion space. To register the DKT segmentation label to native diffusion space, a number of image deformation steps were executed. The DKT segmentation label was moved from MNI space to the PING template space and then inversely warped to subject-specific T1w native space. Using the transformation affine matrix and warp image from *Tractoflow* processing, the DKT label was then aligned from T1w to native diffusion space. This native-space label was generated for each diffusion acquisition and used as input with the corresponding tractogram in *SCIL-scil compute robust connectivity matrix*.*py*, which outputs a connectivity matrix as a *npy* file. Depicted in Figure 5, connectivity is represented as the number of streamlines with endpoints within a region of interest (ROI) label as delineated by the DKT label.

**Figure 5:**
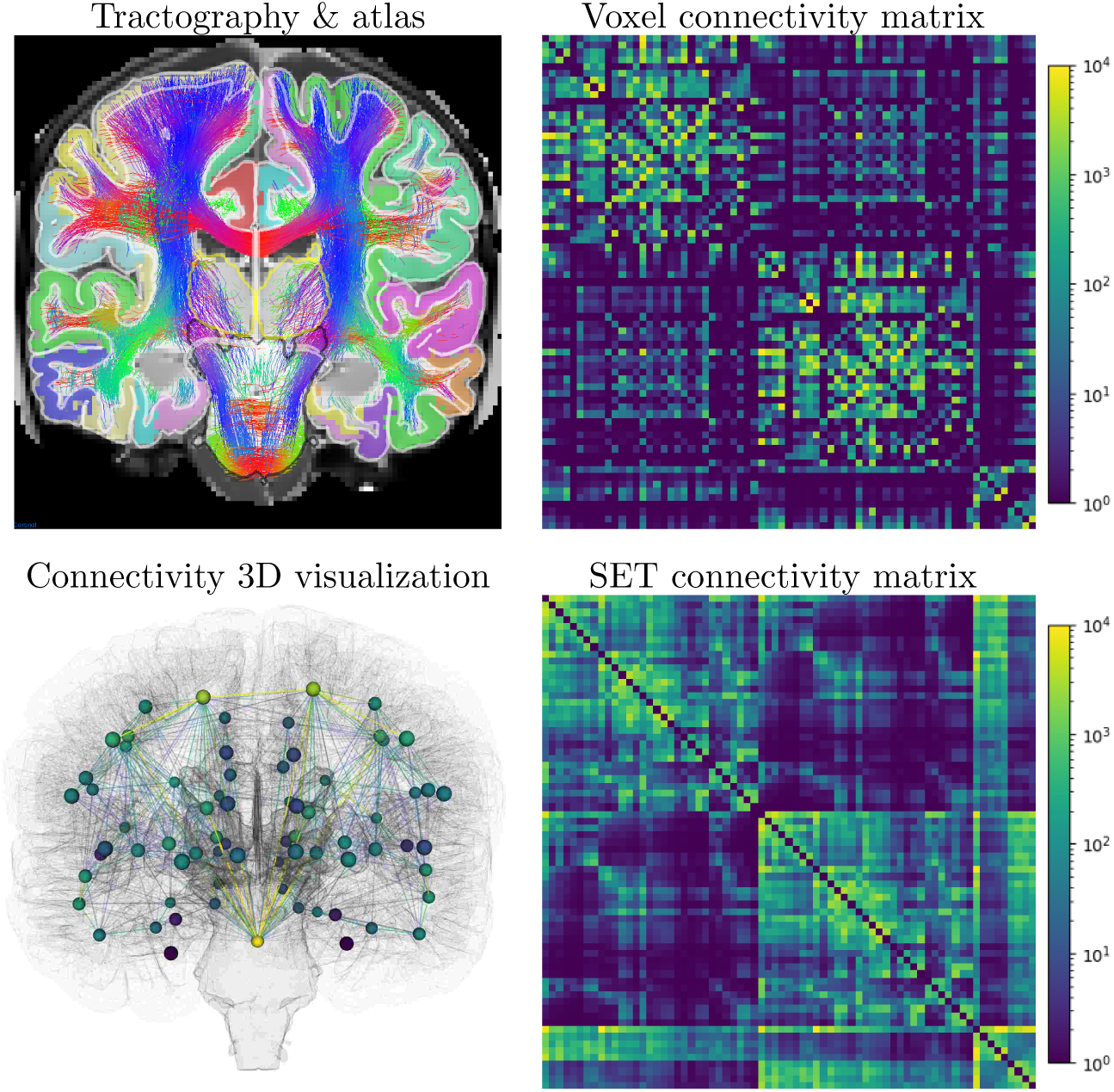
Tractography reconstruction visualized over the DKT atlas labels and resulting connectivity matrices, streamlines count per million in log scale. The connectivity matrix contains 35 cortical and subcortical labels per hemisphere (left, right) and the brainstem, for a total of 71 *×* 71 potential connections.

Using the surface-based method, every streamline generated by SET starts and terminates within a triangle that makes up the cortical surface mesh. Each triangular facet corresponds to a specific cortical label in the CIVET DKT atlas. Subsequently, the connectivity matrix is computed by counting the number of streamlines connecting two different cortical regions.

## 3. Technical Validation

### 3.1. Quality Control

#### Data formatting

Once converted to NIfTI and enforced to RAS orientation, each image header was verified automatically with *MRtrix-mrinfo*, to correct stride convention and data formatting.

#### Anatomical images

If a subject had multiple anatomical T1 acquisitions, a single best-quality image was chosen for each subject, validated visually. During initial assessment of the raw T1 acquisitions, subjects were excluded for severe motion, slicing artifacts, or missing data.

#### CIVET

Brain masks, tissue maps and surfaces from CIVET 2.1 were inspected with the given alignment QC image. Subjects were excluded from further analysis post-CIVET processing due to a) failing to complete processing or b) motion artifacts and incorrectly assigned data.

#### Eddy & Topup

Images were visually assessed before and after Eddy & Topup correction to ensure the correction was applied properly. This also validated that the b0 preliminary mask was not too restrictive.

#### Pre-processed dMRI and T1 registration

Registered pre-processed T1s (masked, denoised, normalized, resampled) were assessed with the fully pre-processed dMRI (prelim masked, denoised, eddy-plus topup-corrected, normalized, resampled) to ensure that the images were aligned and properly pre-processed. In addition, anatomical tissue segmentation maps (WM-GM-CSF) were visually inspected in the diffusion space prior to tractography.

#### Template alignment

Anatomical T1 images and tissue map alignment were verified both qualitatively and quantitatively with an alignment score. Quantitative scores are shown in the next sub-section.

#### DTI & FOD

Diffusion orientations acquired from the b-vector files and their respective eddy- and topup-corrected versions were validated by visualizing DTI main orientation in well-known brain structures: the corpus callosum and the cingulum. A flip in the x axis for b-vector values was required as the images were acquired in LAS orientation, but *Dipy* is based on RAS reference. Resulting local reconstructions were also qualitatively verified for shape, size and orientation discrepancies (Figure 2). This ensured that computed DTI and FOD values were correct and well-aligned with the white matter structure.

#### Diffusion measures

Diffusion measures were also qualitatively validated using maps for FA, MD, RD, RGB and AFD. The variability of these dMRI metrics for this dataset is presented in the next sub-section. Subjects were removed if their diffusion maps presented visible artifacts such as motion, poor registration, uncorrected deformations or signal loss.

#### Tractograms

The resulting tractograms, from 100 randomly chosen subjects, were inspected with 3D visualization, to ensure no reconstruction artifacts were present.

### 3.2. Dataset Variability

In addition to standard quality control and visual validation, quantitative measures and variability were computed after each of the following processing steps. This was done to ensure that no major outliers remained in the final dataset. These variability results could also serve as references for future brain development studies or comparison with the PING dataset.

#### Template alignment

After the template reconstruction (Figure 4), an alignment score (dice) to this averaged template was computed for each individual probabilistic tissue map (Figure 6a). This template alignment score (mean; standard deviation) for WM (0.897; 0.010), GM (0.877; 0.015) and SC (0.930; 0.019) is excellent, with small variance. The CSF score (0.574; 0.039) is lower, compared to others, mostly due to brain mask variation. Subjects misaligned to the template were easily detected as outliers from the WM dice score (z-score *<* -5) and confirmed through visual validation. In general, a data point is considered an outlier when the z-score is greater than 3 or less than -3. Removing these non-aligned subjects consistently reduced the dice standard deviation by a factor of 3 for WM (around 2 for other tissue maps). Almost identical results and corresponding outliers were found using other distance measures (L1 and L2) as well.

**Figure 6:**
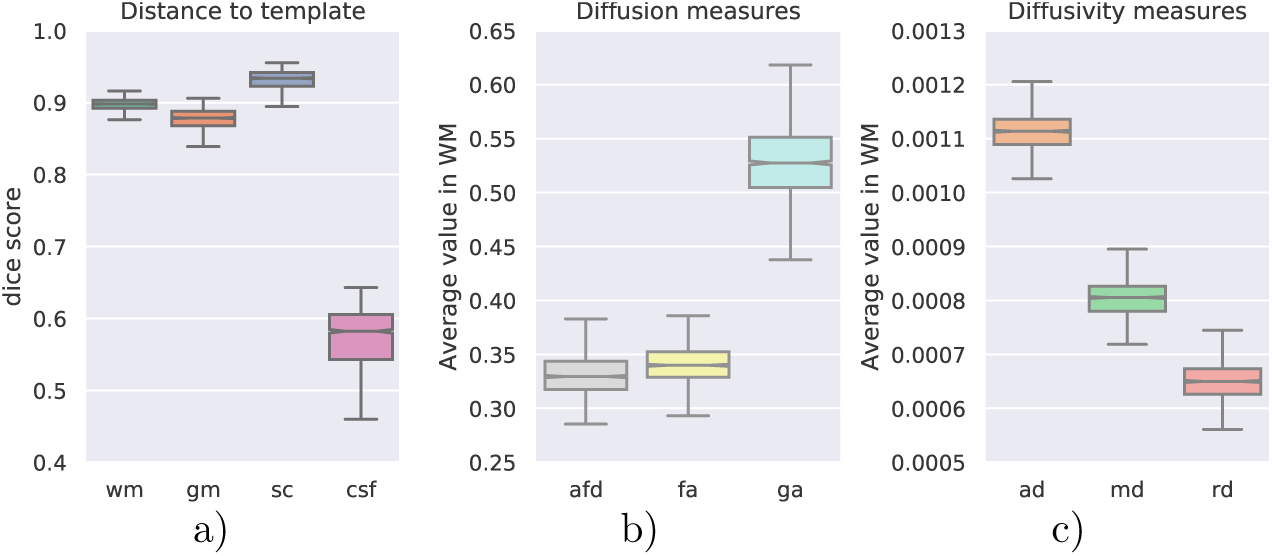
a) Dice alignment score to the template, for each subject. White matter measures of: b) diffusion (AFD, FA, GA) and c) diffusivity in *mm*^2^*/s* (AD, MD, RD).

#### Diffusion measures

Measures estimated from dMRI models (DTI and FOD) were also evaluated for each subject. Intensities in each tissue structure were analysed with a histogram and average value per subject (see Figure 6, Table A.1). From this variability analysis, results suggest that some FOD metrics, such as NUFO, are too variable to measure any group differences in this dataset. Nonetheless, this limitation could be partially caused by the variable nature of this dataset (age variation, multiple acquisition site, etc).

#### FOD model data response (frf)

Fiber response function (frf) was first estimated for each dMRI acquisition in the center of the WM structure in high FA values near the corpus callosum. Results were then compared for each scanner and acquisition site (Figure 7a, Table A.2). Since all frf were similar across PING, values used to compute FOD were fixed to 18, 4, 4 (ratio of 18*/*4), the dataset average.

**Figure 7:**
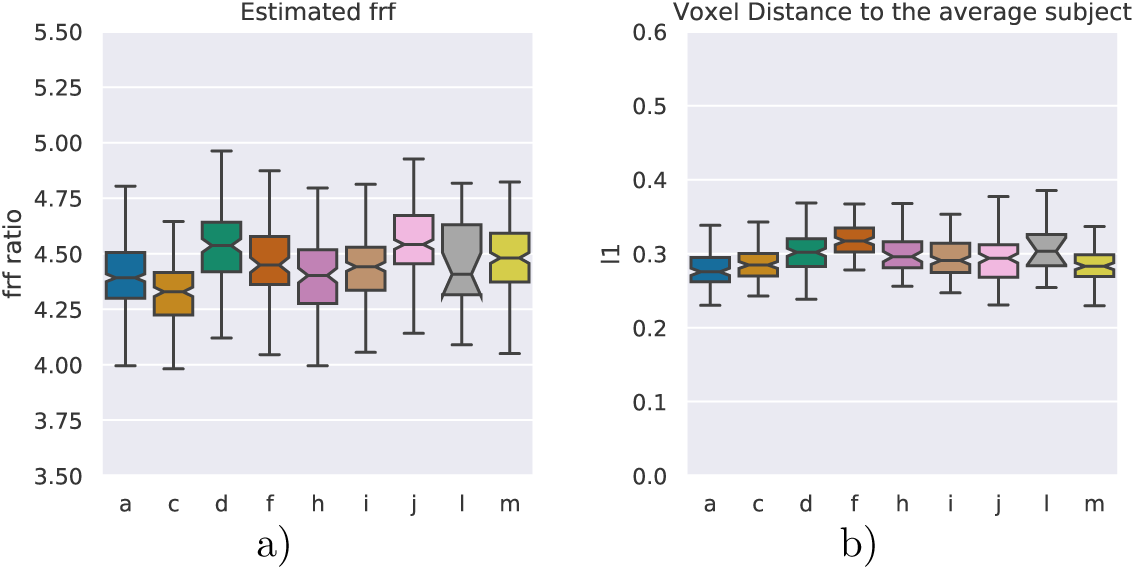
a) Estimated fiber response function (frf) ratio, b) connectivity matrices distance to the average subject, across acquisition sites. MRI scanner for each site: Philips Achieva (a,h), GE Signa HDx (c), GE Discovery MR750 (i) and Siemens TrioTim (d,f,j,l,m).

#### Connectivity matrices

Connectivity matrices were employed to measure inter-subject distance. For subjects with multiple dMRI acquisitions, intra-subject distance was also measured. In addition, variability analysis was performed over connectivity matrices to estimate outliers and the differences between acquisition sites and age groups. Figure 7b displays L1-distance, from the average connectivity to each subject’s connectivity matrix, grouped per acquisition site (Table A.3).

#### Metrics

Dice score and distance equations, in between images *A* and *B* at each voxel position *v* in the whole 3D volume 𝒱 is as follows:

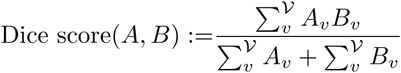

## 4. Conclusion

In this work, we created a processed and quality-assessed set of connectivity derivatives of the PING dataset, using a robust dMRI processing pipeline, *Tractoflow* [Theaud et al., 2020]. Resulting derivatives were assessed and validated at multiple steps to ensure a high-quality, normative dataset. Using a state-of-the-art pipeline and openly publishing the dataset will encourage the use of dMRI metrics within the scope of PING-related connectomic analyses. The resulting dataset includes:

- processed T1 and diffusion weighted images, from established software,
- brain tissue segmentation maps along with cortical surfaces from CIVET,
- advanced dMRI processing using fiber orientation distribution (FOD) in addition to diffusion tensor imaging (DTI),
- WM pathways reconstruction from cutting edge tractography algorithms,
- structural connectivity matrices, estimated using voxel and surface approach.

### Code availability

An adapted version of *Tractoflow* for PING processing is available in the complementary files on NDA, named *Tractoflow-pve*. Within this project folder, both `‘USAGE’` and `‘README.md’` give information about usage, `‘nextflow.config’` contains configuration details, and `‘main.nf’` includes all processing steps and details. The singularity image `‘tractoflow_2.0.0.img’` contains all requirements to execute the `‘tractoflow-pve/’` pipeline. A secondary sub-pipeline `‘set-nf/’` is provided to compute the “surface-enhanced tractography” and surface-based measures, using the same file organisation as in `‘tractoflow-pve’` with the addition of `‘set_ping.img’` to execute it. The `‘set_ping.img’` is also used to execute the `‘civet-nf’` pipeline which organizes PVE output files from CIVET 2.1 for `‘tractoflow-pve’`. All code is packaged within the `‘Processing_PING’` archive file available on NDA.

All dMRI and T1w processing software employed in this work are available online and free of use for researchers: *ANTs, CIVET, dcm2niix, FreeSurfer, FSL, MRtrix, Nibabel*, and *Dipy*. Processing tools utilized for scripts and pipelining: *Nextflow, Singularity*, and *Python*. Visualisation tools: *Fury, MI-Brain, MisterI*, and *MRtrix mrview*.

### Data Records

Data and research tools used in the preparation of this manuscript were obtained from the National Institute of Mental Health (NIMH) Data Archive (NDA). NDA is a collaborative informatics system created by the National Institutes of Health to provide a national resource to support and accelerate research in mental health. Dataset identifier(s): 10.15154/1519178. This manuscript reflects the views of the authors and may not reflect the opinions or views of the NIH or of the Submitters submitting original data to NDA. Information on the original PING dataset, including acquisition protocols, is detailed in Jernigan et al. [2016].

The processed derivatives described here and the code used to produce them are all available on the NDA website, study ID #932 titled *Diffusion-*weighted derivatives of PING, doi: 10.15154/1519178. All code and file descriptions are found at the study’s page under ‘Data Analysis,’ and resulting derivative data are packaged in the ‘Results’ section. Descriptions for all derivative files and in which data package they are organized can be found in *ping dwiderivatives ndarinfo*.*xlxs* under ‘Data Analysis.’

### Processed anatomical maps

All anatomical 3D volumes and derived maps are saved in *NIfTI* format, supported in *dcm2niix, ANTs, Nibabel*, and *MRtrix*.

### Processed dMRI

Images at each processing step are saved in 3D or 4D (representing a series of 3D images) *NIfTI* : b0 volumes (A-P, P-A), dMRI directions, as well as upsampled denoised-, eddy- & topup-corrected images.

### Diffusion bval & bvec

Eddy-corrected b-values and b-vectors are saved in the same format as initially converted by *dcm2niix* : b-values are listed in a text file (space-separated), b-vector directions are listed in x-y-z format in distinct rows.

### Cortical surfaces

The Visualization Toolkit (VTK) format was used for all meshes, with vertices saved in world coordinates (LPS). In this format, cortical surfaces can be visualized and overlaid onto the T1 image with *MI-Brain* (see Figure 5).

### DTI

Tensor images were saved with *Dipy* and *Nibabel* in NIfTI format. The resulting image is a 4D volume: spatially represented in 3D with a fourth dimension containing all 6 symmetric tensor coefficients. Tensor coefficients were saved in this given order along the fourth dimension axis, where: [*D*_00_, *D*_01_, *D*_02_, *D*_11_, *D*_12_, *D*_22_], where

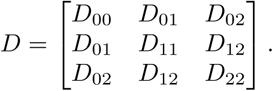

FOD fields were saved with *Dipy* using the default spherical harmonics (SH) representation [Descoteaux et al., 2009]. Only even-ordered SH coefficients 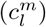 were saved, as computed FOD are symmetric. The SH file is ordered by SH order (*l*) first and phase (*m*) second: 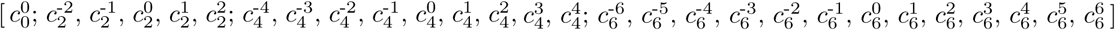.

#### Tractograms

Tractography reconstructions were saved in *TrackVis* format (`‘.trk’`). This format is well supported in *Nibabel, Dipy* and *MI-Brain*.

#### Connectivity matrices

All resulting connectivity matrices were stored as nummpy array matrices (`‘.npy’`) format. The ordered list of labels is given in the `‘labels_co_matrix.txt’` file.

### Usage Notes

#### Command lines

*Nextflow* and *Singularity* need to be installed; a complete example is given in `‘PING_example`.`sh’`.

~~~
nextflow run civet-nf/main.nf --civet path/to/civet·output/ “
   -with-singularity set ping.img
sh civet-nf/tree·for·tractoflow.sh -f path/to/data/ -c civet-nf/results/ “
   -o tractoflow·input/
nextflow run tractoflow-pve/main.nf --root tractoflow·input/ “
   --dti·shells “0 1000” --fodf·shells “0 1000” “
   -profile civet·pve   -with-singularity tractoflow·2.0.0.img
nextflow run set-nf/main.nf --tractoflow tractoflow·nf/results/ “
   --civet path/to/civet·output/ -profile civet·DKT “
   -with-singularity set·ping.img
~~~

#### Pediatric Imaging Neurocognition and Genetics (PING)

Data collection and subsequent dataset for this project were obtained from the Pediatric Imaging, Neurocognition and Genetics Study (PING), National Institutes of Health Grant RC2DA029475. PING is funded by the National Institute on Drug Abuse and the Eunice Kennedy Shriver National Institute of Child Health & Human Development. PING data are disseminated by the PING Coordinating Center at the Center for Human Development, University of California, San Diego. PING data (nda.nih.gov/editcollection.html?id=2607) and Consortium (ping-dataportal.ucsd.edu/sharing/Authors10222012.pdf), detailed in Jernigan et al. [2016].

## Acknowledgements

Supported by grants from Brain Canada (238990, 243030), CFREF/HBHL Innovative Ideas (247613), Coutu Research Fund (241177), CFREF/HBHL Discovery (247712), Université de Sherbrooke Research Chair in NeuroInformatics and NSERC Discovery Grant.

## Author contributions statement

The concept of this work, processing the structural connectivity of the PING dataset, was proposed and realized by Noor B Al-Sharif, supervised by Alan C Evans. Diffusion MRI processing planning and work outline was done by Etienne St-Onge, supervised by Maxime Descoteaux. *Tractoflow* pipeline initial implementation, and technical assistance was done by Guillaume Theaud. Processing pipeline adaptation for CIVET tissue maps, surfaces and SET was implemented by Etienne St-Onge, and rigorously tested by Noor B Al-Sharif, resulting in *Tractoflow-pve*.

## Data Citations

Data and research tools used in the preparation of this manuscript were obtained from the National Institute of Mental Health (NIMH) Data Archive (NDA). NDA is a collaborative informatics system created by the National Institutes of Health to provide a national resource to support and accelerate research in mental health. Dataset identifier(s): 10.15154/1519178. This manuscript reflects the views of the authors and may not reflect the opinions or views of the NIH or of the Submitters submitting original data to NDA.

## Appendix A. Full Comparison

**Table A.1:** Diffusivity measures (AD, MD, RD) in *mm*^2^*/s*, diffusion measures (AFD, FA, GA) and tissue maps dice alignment score, average & variance over all subjects, for each segmentation maps.

|  | WM | GM | SC | CSF |
| --- | --- | --- | --- | --- |
| AD | 0.00111, 0.000033 | 0.00116, 0.000072 | 0.00104, 0.000033 | 0.00160, 0.000146 |
| MD | 0.00080, 0.000033 | 0.00103, 0.000059 | 0.00082, 0.000032 | 0.00144, 0.000125 |
| RD | 0.00065, 0.000035 | 0.00096, 0.000054 | 0.00070, 0.000033 | 0.00135, 0.000117 |
| AFD | 0.332, 0.022 | 0.332, 0.026 | 0.360, 0.027 | 0.244, 0.030 |
| FA | 0.340, 0.019 | 0.129, 0.011 | 0.254, 0.015 | 0.114, 0.020 |
| GA | 0.529, 0.038 | 0.191, 0.026 | 0.375, 0.024 | 0.190, 0.050 |
| dice | 0.897, 0.010 | 0.877, 0.015 | 0.930, 0.019 | 0.574, 0.039 |

**Table A.2:** Estimated fiber response function (frf) in *mm*^2^*/s* across acquisition sites: first eigenvalue, second eigenvalue, ratio. Philips Achieva (a,h), GE Signa HDx (c), GE Discovery MR750 (i) and Siemens TrioTim (d,f,j,l,m).

|  | first eigenvalue | second eigenvalue | ratio |
| --- | --- | --- | --- |
| a | 0.001709, 0.000124 | 0.000388, 0.000026 | 4.40, 0.16 |
| c | 0.001686, 0.000076 | 0.000391, 0.000019 | 4.32, 0.14 |
| d | 0.001887, 0.000196 | 0.000416, 0.000041 | 4.54, 0.18 |
| f | 0.001850, 0.000207 | 0.000413, 0.000041 | 4.47, 0.17 |
| h | 0.001836, 0.000201 | 0.000418, 0.000039 | 4.39, 0.18 |
| i | 0.001716, 0.000087 | 0.000387, 0.000019 | 4.44, 0.16 |
| j | 0.001932, 0.000211 | 0.000425, 0.000042 | 4.54, 0.18 |
| l | 0.001735, 0.000128 | 0.000390, 0.000022 | 4.45, 0.18 |
| m | 0.001782, 0.000183 | 0.000397, 0.000036 | 4.49, 0.19 |

**Table A.3:** Connectivity matrices L1, L2 and *χ*^2^ distances, per acquisition site, to the average matrix. Philips Achieva (a,h), GE Signa HDx (c), GE Discovery MR750 (i) and Siemens TrioTim (d,f,j,l,m).

|  | voxel L1 | voxel L2 | SET L1 | SET L2 |
| --- | --- | --- | --- | --- |
| a | 0.278, 0.027 | 0.0234, 0.0029 | 0.385, 0.040 | 0.0331, 0.0055 |
| c | 0.287, 0.026 | 0.0242, 0.0032 | 0.382, 0.036 | 0.0303, 0.0068 |
| d | 0.301, 0.028 | 0.0254, 0.0030 | 0.375, 0.041 | 0.0278, 0.0050 |
| f | 0.317, 0.024 | 0.0266, 0.0031 | 0.417, 0.041 | 0.0331, 0.0058 |
| h | 0.299, 0.026 | 0.0251, 0.0029 | 0.435, 0.037 | 0.0370, 0.0048 |
| i | 0.291, 0.024 | 0.0245, 0.0028 | 0.358, 0.038 | 0.0269, 0.0051 |
| j | 0.291, 0.033 | 0.0243, 0.0033 | 0.378, 0.044 | 0.0292, 0.0057 |
| l | 0.310, 0.031 | 0.0252, 0.0027 | 0.429, 0.038 | 0.0347, 0.0063 |
| m | 0.283, 0.023 | 0.0236, 0.0026 | 0.363, 0.039 | 0.0274, 0.0052 |

